## SupplementaryData for "Histone sample preparation for bottom-up mass spectrometry: a roadmap to informed decisions"

Supplementary data

**Supplementary Figure 1.** Coverage plots of all core histones from the AQUA heavy histone peptide standard from Li et al [15].

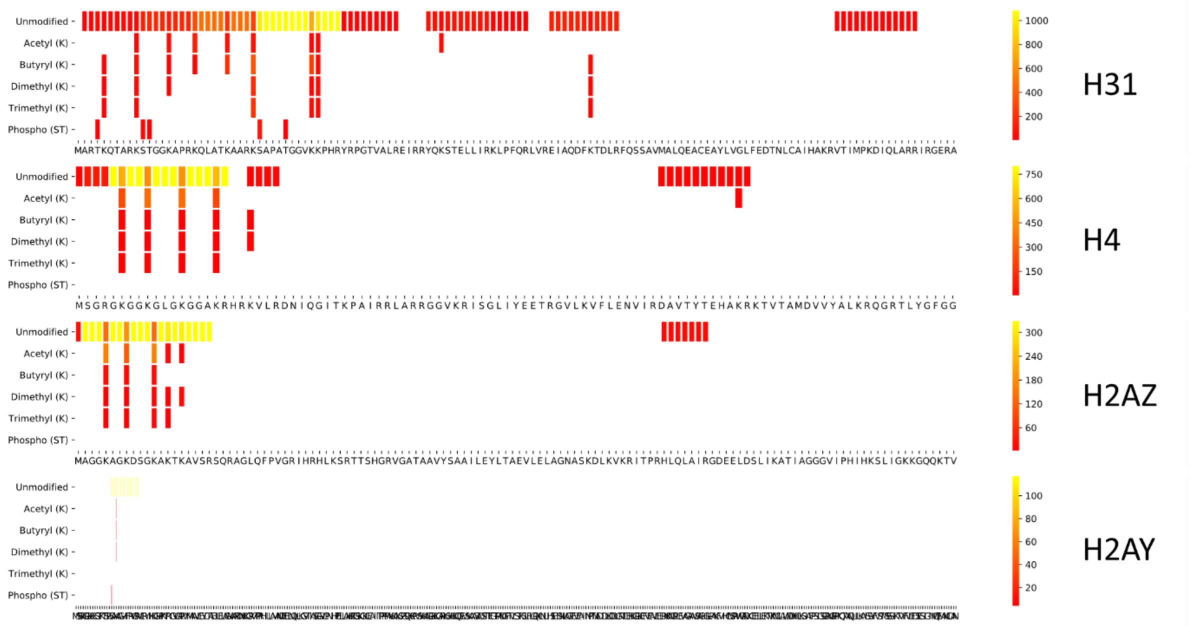


**Supplementary Figure 2.** Mascot example of peptide H2B 1-29 with multiple possible annotations above scoring threshold for the same mass spectrum. The first hit is reported in the spectrum with a score of 63.4 and expectancy value of 4.6 e-6.

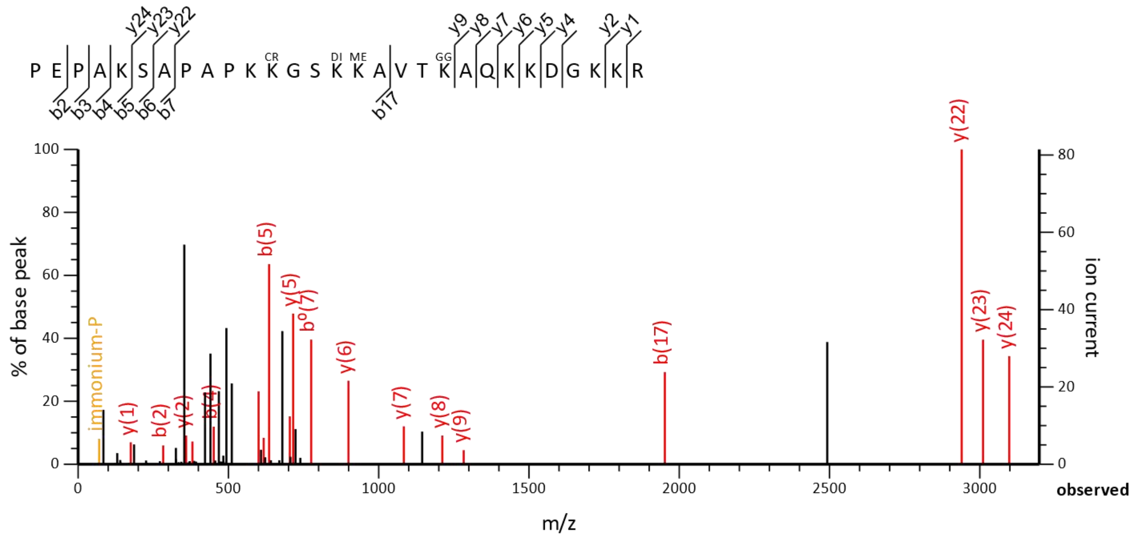


| Score | Mr(calc) | Delta | Sequence | Site Analysis |
|---|---|---|---|---|
| 63.4 | 3731.1350 | -0.0253 | PEPAKSAPAPKKGSKKAVTKAQKKDGKKR | Methyl KR K16, Crotonyl K12, GG Prop K20, Dimethyl KR K15; 0.02% |
| 63.4 | 3731.0986 | 0.0111 | PEPAKSAPAPKKGSKKAVTKAQKKDGKKR | |
| 63.4 | 3731.1350 | -0.0253 | PEPAKSAPAPKKGSKKAVTKAQKKDGKKR | Methyl KR K11, Crotonyl K12, GG Prop K20, Dimethyl KR K15; 0.02% |
| 60.4 | 3731.1350 | -0.0253 | PEPAKSAPAPKKGSKKAVTKAQKKDGKKR | Methyl KR K20, Crotonyl K12, GG Prop K16, Dimethyl KR K15; 0.01% |
| 60.4 | 3731.0986 | 0.0111 | PEPAKSAPAPKKGSKKAVTKAQKKDGKKR | |
| 60.4 | 3731.1350 | -0.0253 | PEPAKSAPAPKKGSKKAVTKAQKKDGKKR | |
| 60.4 | 3731.1350 | -0.0253 | PEPAKSAPAPKKGSKKAVTKAQKKDGKKR | |
| 60.4 | 3731.1602 | -0.0505 | PEPAKSAPAPKKGSKKAVTKAQKKDGKKR | |
| 60.4 | 3731.1350 | -0.0253 | PEPAKSAPAPKKGSKKAVTKAQKKDGKKR | |
| 60.4 | 3731.1350 | -0.0253 | PEPAKSAPAPKKGSKKAVTKAQKKDGKKR | Methyl KR K20, Crotonyl K12, GG Prop K11, Dimethyl KR K15; 0.01% |

**Supplementary Figure 3.** Coverage plot of Histone H3 for each workflow highlighting the chemical noise and obscured regions introduced by PropTryp.

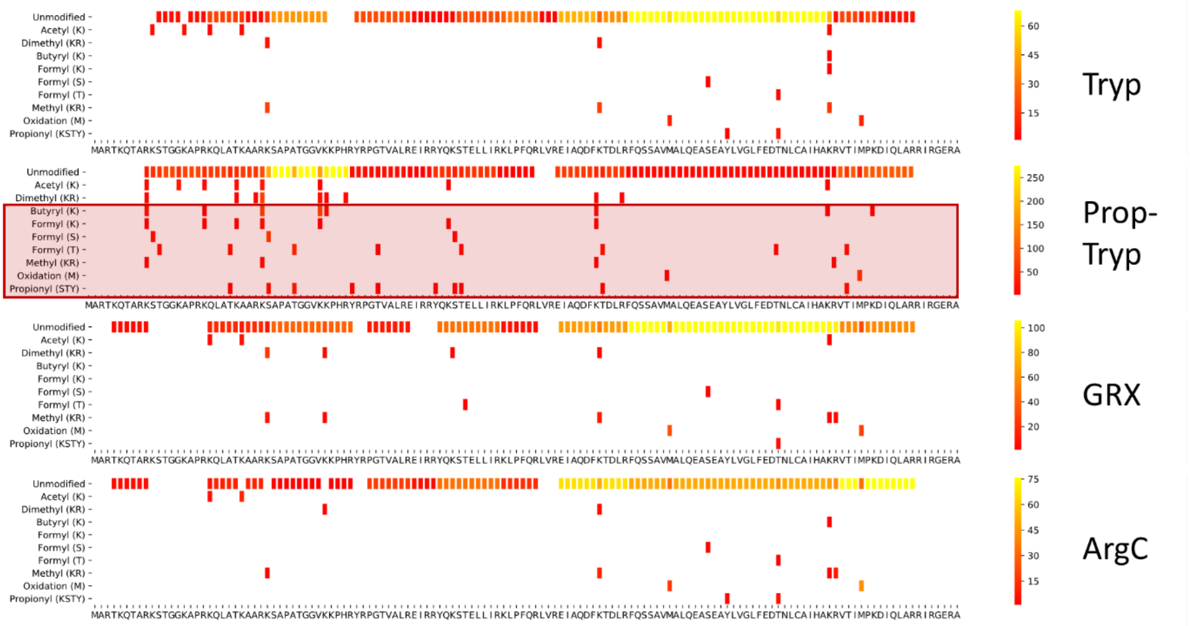


**Supplementary Figure 4.** Coverage plot of Histone H4 for each workflow highlighting the chemical noise and obscured regions introduced by PropTryp.

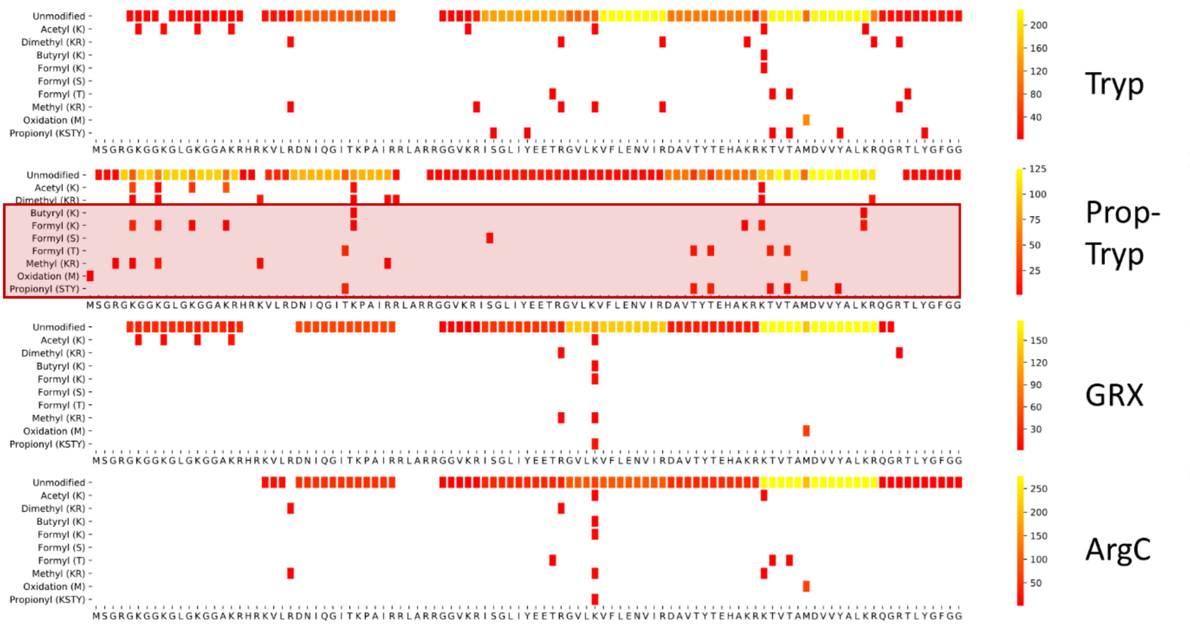


**Supplementary Figure 5.** Coverage plot of Histone H2A for each workflow highlighting the chemical noise and obscured regions introduced by PropTryp.

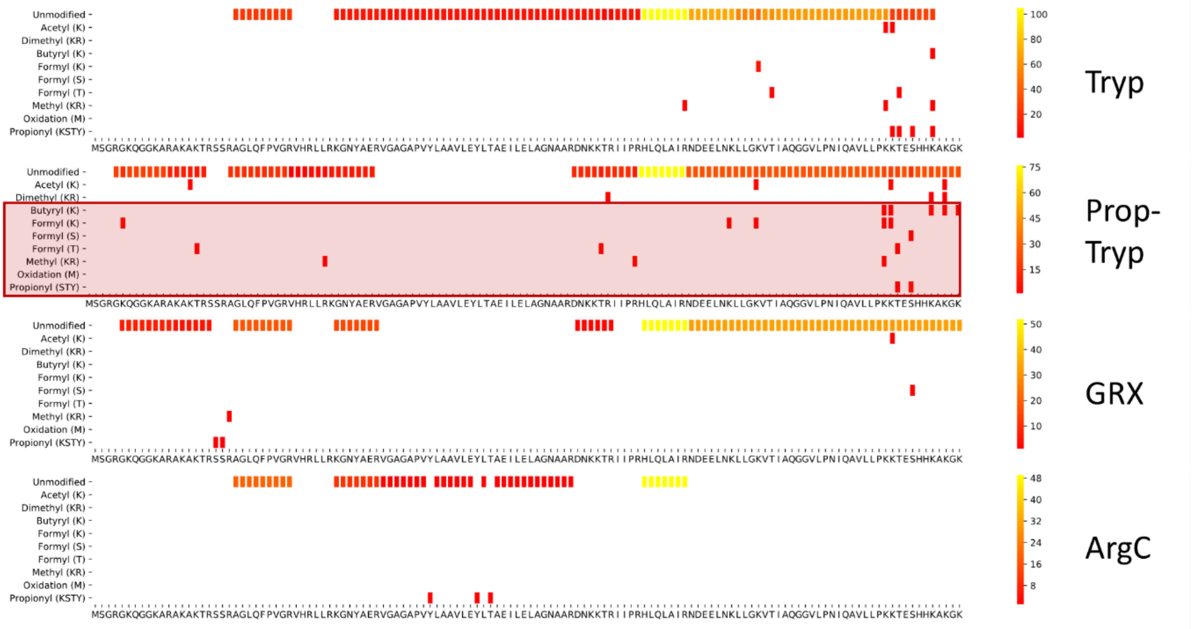


**Supplementary Figure 6.** Coverage plot of Histone H2B for each workflow highlighting the chemical noise and obscured regions introduced by PropTryp.

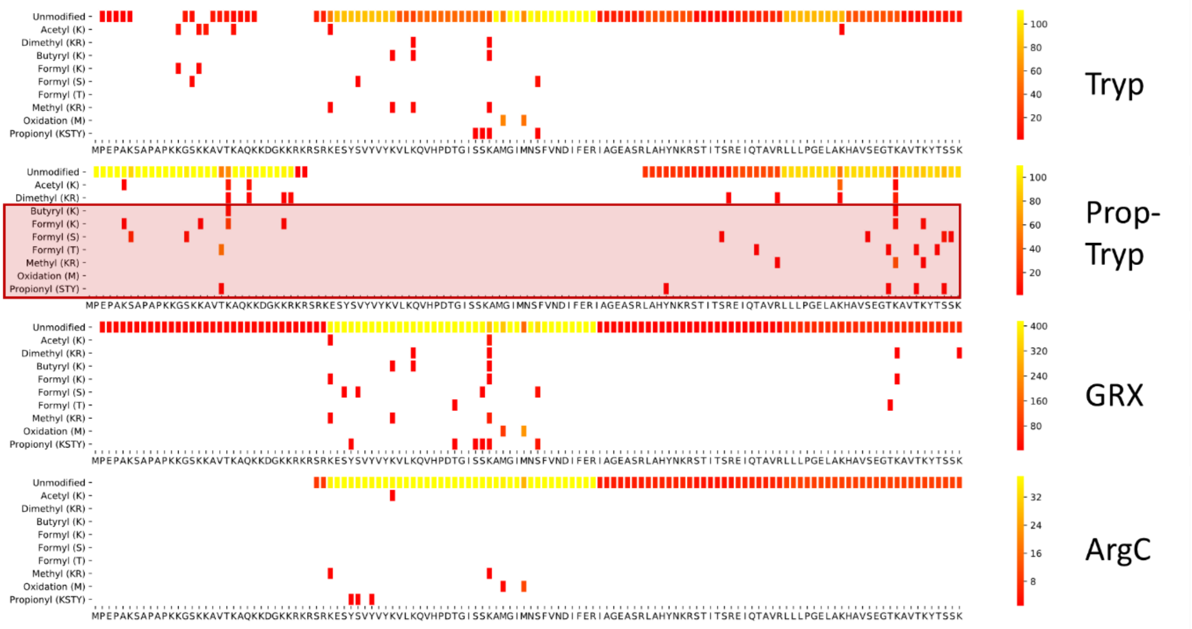


**Supplementary Figure 7**. Coverage plot of Histone H4 for each workflow with the predicted coverage derived from Uniprot as a reference.

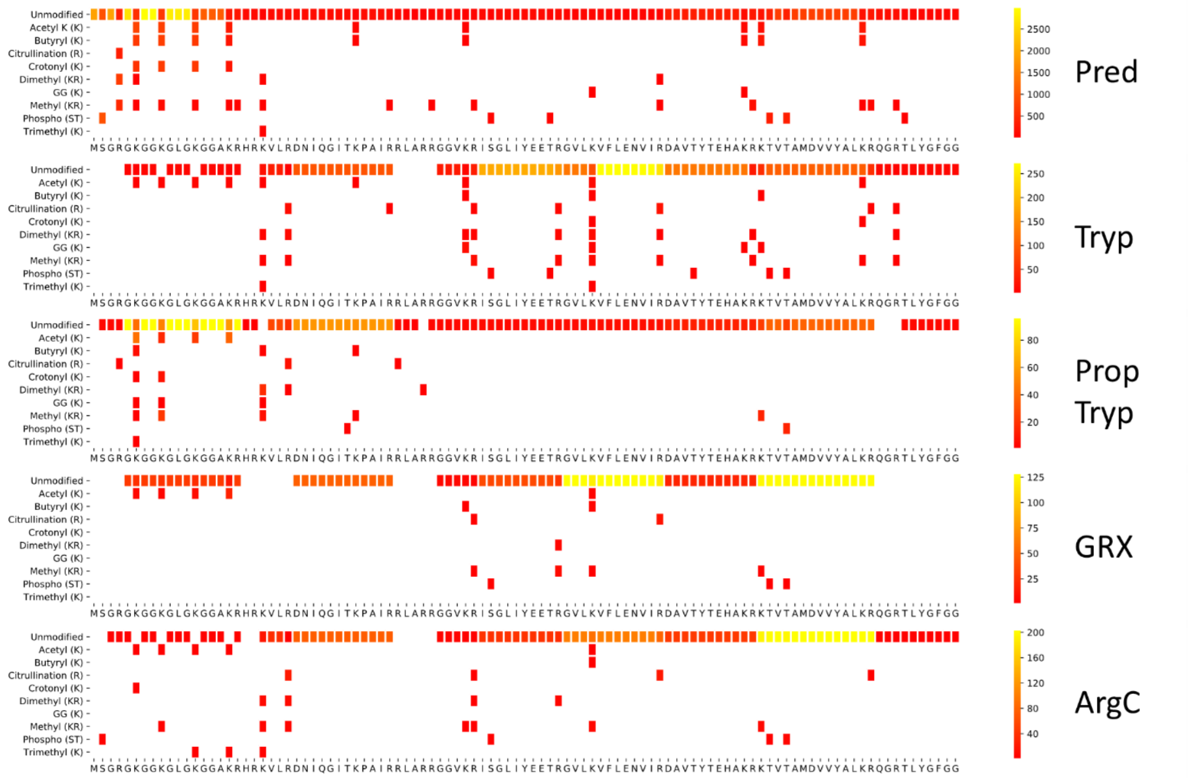


**Supplementary Figure 8**. Coverage plot of Histone H2A for each workflow with the predicted coverage derived from Uniprot as a reference.

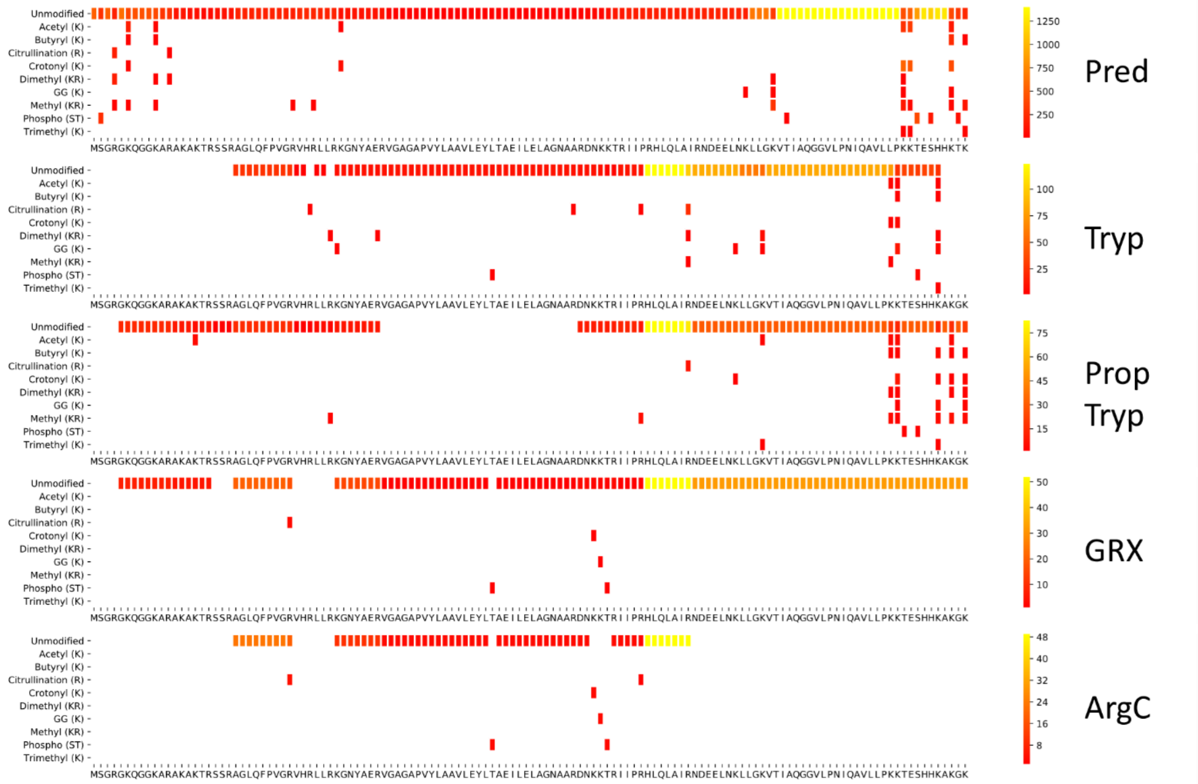


**Supplementary Figure 9**. Coverage plot of Histone H2B for each workflow with the predicted coverage derived from Uniprot as a reference.

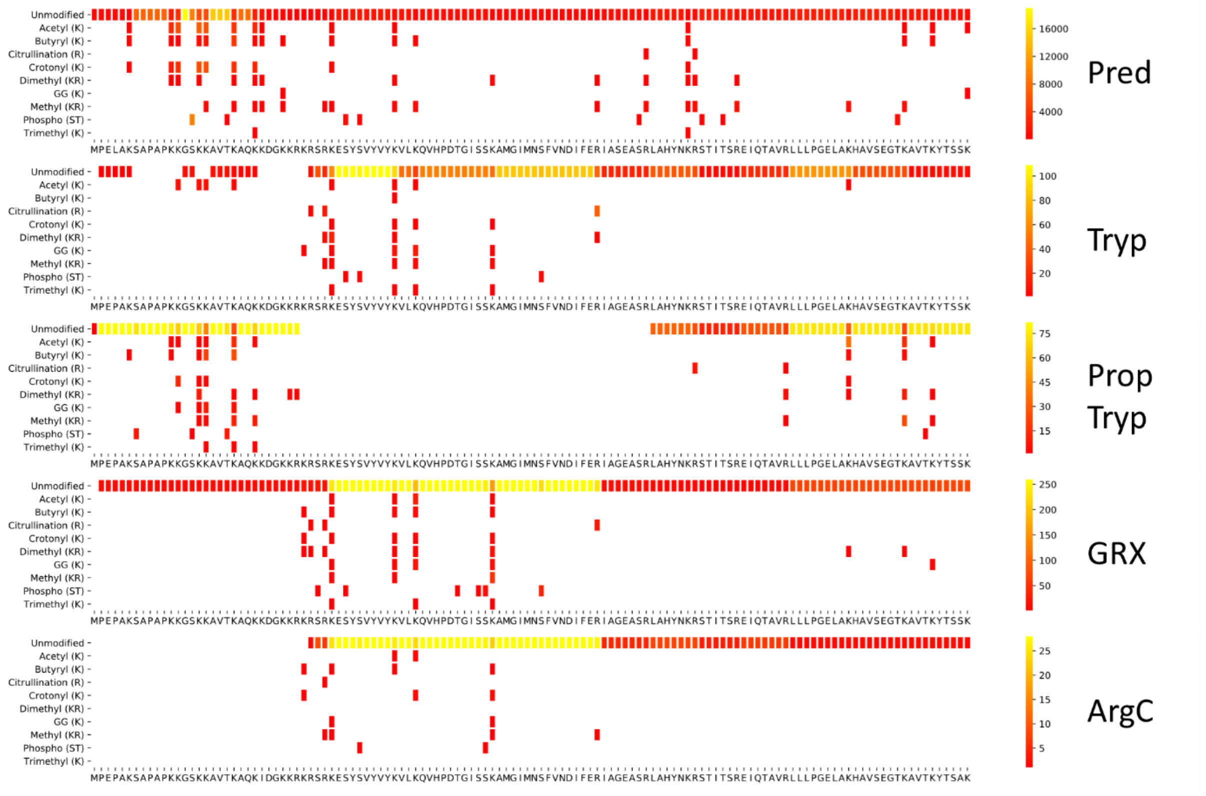


**Supplementary Figure 10**. Coverage plot of Histone H1 for each workflow with the predicted coverage derived from Uniprot as a reference. The ArgC workflow did not cover Histone H1.

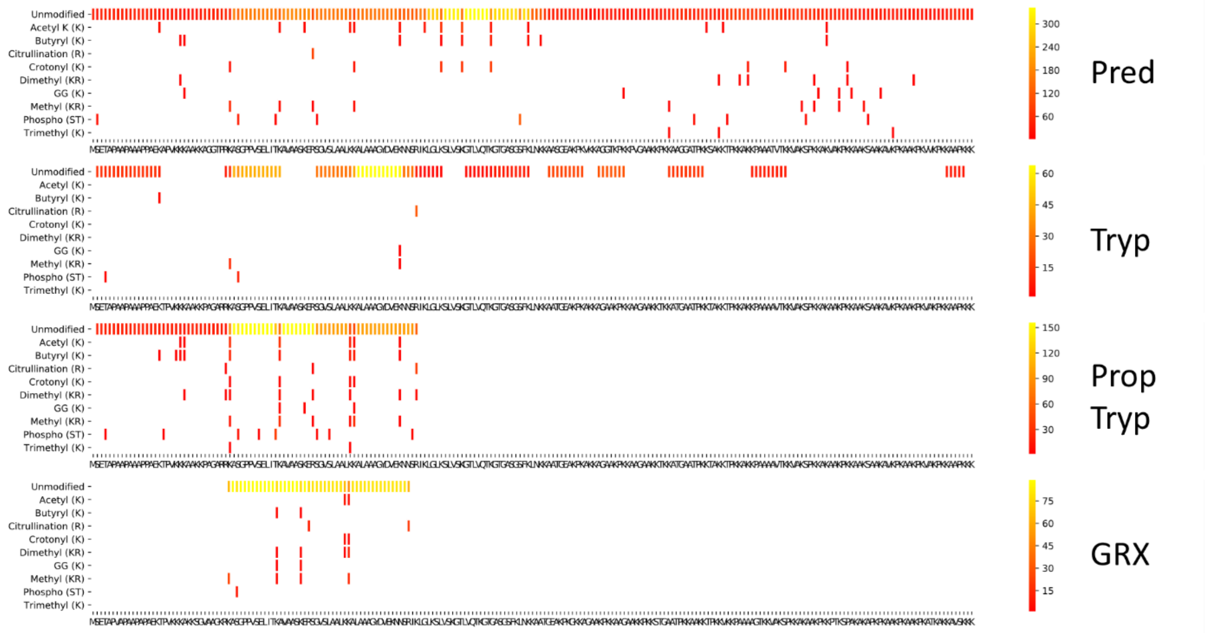


**Supplementary Figure 11: feature detectability for different enzymatic treatments**. **(A-C)** Two dimensional consensus representations of the different treatments with retention time in the y-axis from top to bottom and the m/z on the X-Axis. Note that the retention time is representated by scan rate and is not linear. Note that the spread of features across the LC gradient is better for PropTryp.

**(D)** Charge state distribution of all annotated ions.

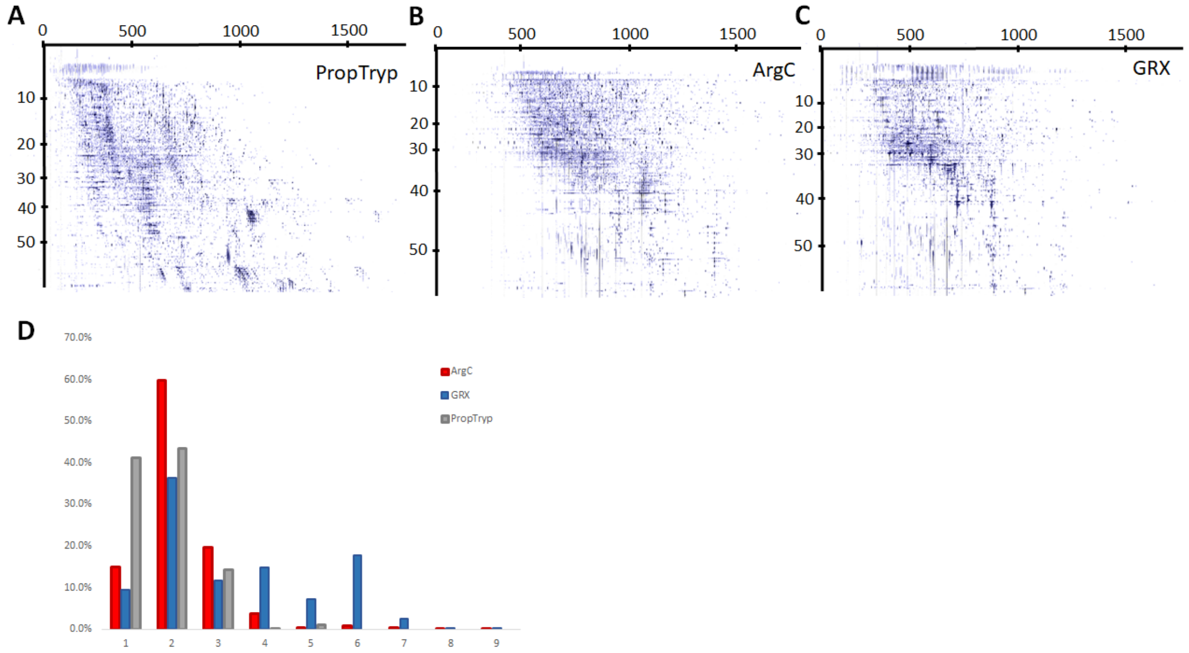


**Supplementary Figure 12**. Coverage plot of the Semi-specific arginine cleavage search on the ArgC workflow data shows a great coverage of all histone proteins and their modifications. Still, enzymatic aspecificity complicates downstream quantification as shown in Figure 7.

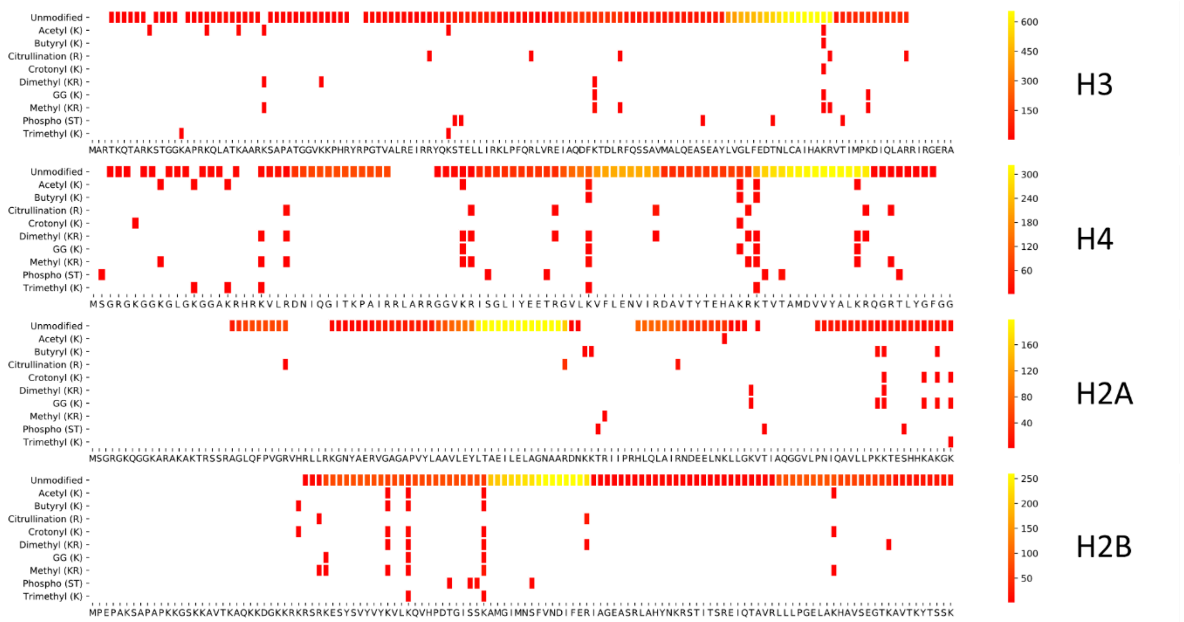


**Supplementary Figure 13.** Localization of the most prominent peptidoforms of the PropTryp workflow in the 3D ion space of the Progenesis QIP software.

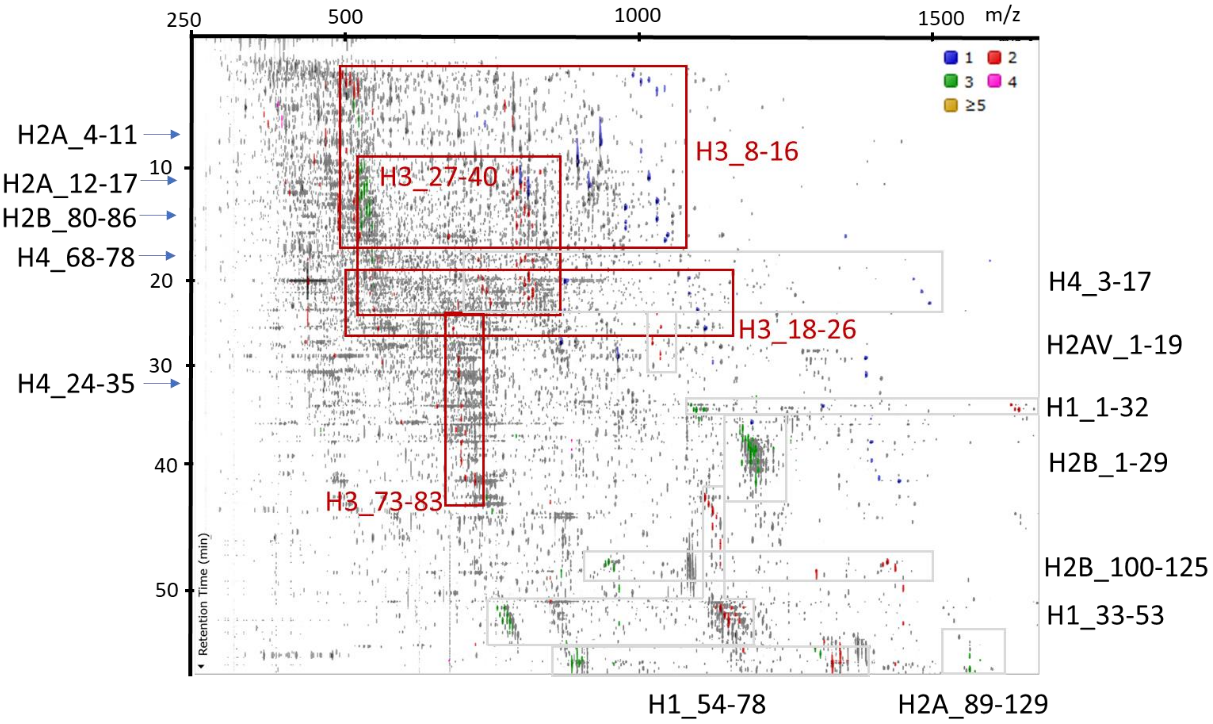


**Supplementary Figure 14.** Localization of the most prominent peptidoforms of the GRX workflow in the 3D ion space of the Progenesis QIP software.

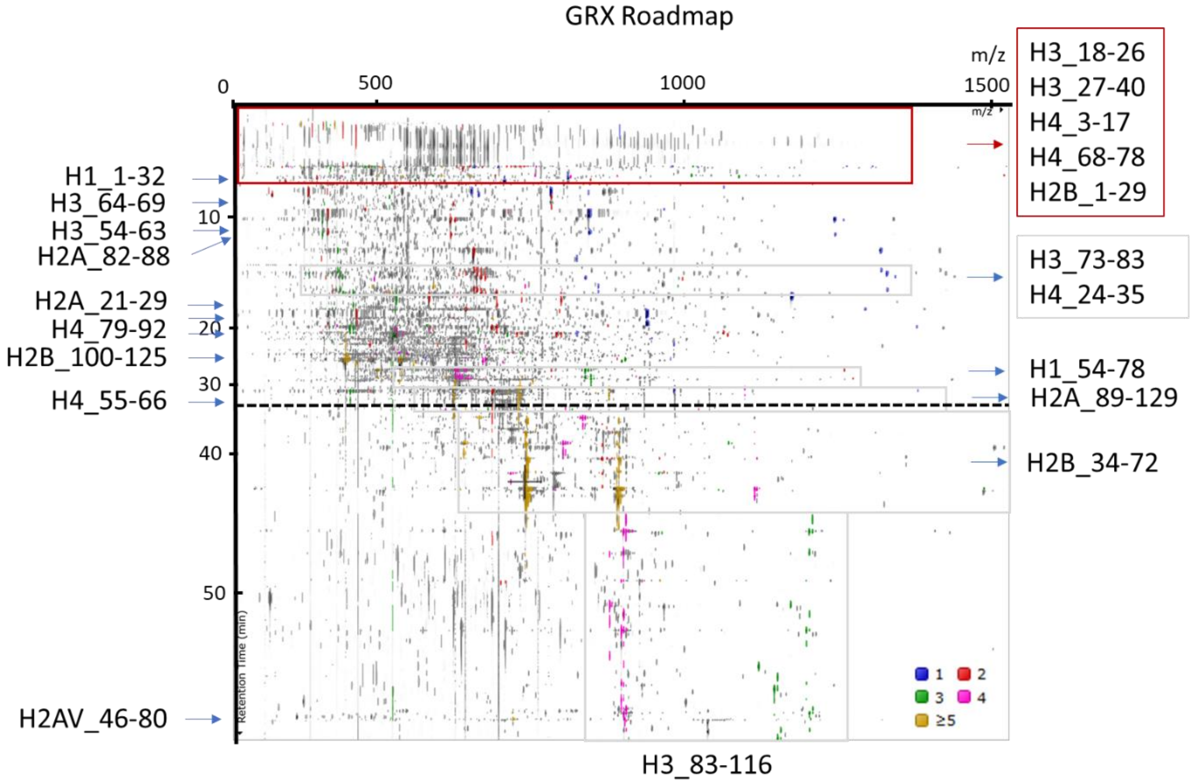


**Supplementary table 1.** All features exported from Progenesis QIP for all workflows. Detailed legend can be found in the excel file

**Supplementary table 2.** All annotated features exported from Progenesis QIP for the ArgC, GRX, and PropTryp workflow. Detailed legend can be found in the excel file.
